# Mean Dimension of Generative Models for Protein Sequences

**DOI:** 10.1101/2022.12.12.520028

**Authors:** Christoph Feinauer, Emanuele Borgonovo

## Abstract

Generative models for protein sequences are important for protein design, mutational effect prediction and structure prediction. In all of these tasks, the introduction of models which include interactions between pairs of positions has had a major impact over the last decade. More recently, many methods going beyond pairwise models have been developed, for example by using neural networks that are in principle able to capture interactions between more than two positions from multiple sequence alignments. However, not much is known about the inter-dependency patterns between positions in these models, and how important higher-order interactions involving more than two positions are for their performance. In this work, we introduce the notion of mean dimension for generative models for protein sequences, which measures the average number of positions involved in interactions when weighted by their contribution to the total variance in log probability of the model. We estimate the mean dimension for different model classes trained on different protein families, relate it to the performance of the models on mutational effect prediction tasks and also trace its evolution during training. The mean dimension is related to the performance of models in biological prediction tasks and can highlight differences between model classes even if their performance in the prediction task is similar. The overall low mean dimension indicates that well-performing models are not necessarily of high complexity and encourages further work in interpreting their performance in biological terms.

## 1 Introduction

Generative models for protein sequences trained on datasets of homologous sequences are used widely for mutational effect prediction [1, 2, 3], pathogenicity prediction in humans [4], the creation of novel protein sequences [5], and structure prediction [6] and protein interaction prediction [7, 8]. While pairwise models that consider the compatibility between pairs of positions when assigning the probability to a sequence have been used and studied extensively in this field [9, 6, 2, 3, 10, 11], the question of how to include higher-order epistasis into these models is a topical research subject. While it is possible to manually add specific higher-order interactions to pairwise models [12], more recent works explore the use of neural networks, which can in principle capture arbitrary patterns from the underlying data. Architectures that have been proposed in literature are based for example on Variational Autoencoders [13, 1], GANs [14] and Transformer based architectures [15].

Nonetheless, it is often unclear what patterns these more complex models capture in the data and whether higher-order interactions play a key role in their performance. This is an important question, since disentangling biologically relevant effects from biases originating in phylogenetic relationships in the training sequence and data preprocessing pipelines might reveal interesting biological insights and lead to improved models. Some recent work has attempted to address this issue, for example by introducing quantities like the *pairwise saliency* and relating it to the performance in structural and mutational effect prediction [16], by extracting pairwise models from the original neural network models and comparing their performance [17], or by assessing how well-trained models reproduce sequence statistics [18].

In this work, we propose to analyze the *mean dimension* of generative protein sequence models, a concept that originates in statistics [19] and has recently been used for the analysis of neural networks trained for supervised prediction tasks [20, 21]. To extend the definition of mean dimension to probabilistic models *p*(*s*) over amino acid sequences *s*, we define it as a weighted average over interaction orders *k*, where every order is weighted by its independent contribution *c_k_* to the variance of log *p*(*s*) when *s* is sampled from the uniform distribution. We show below that, in this setting,

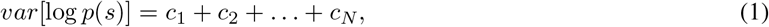

where *N* is the length of the amino acid sequences and *c_k_* depends only on terms in log *p*(*s*) that involve *k* positions in the sequence. The mean dimension *D_f_* can then be defined as

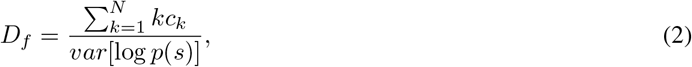

where every interaction order is weighted by their contribution to the variance (see Fig. 1 for an illustration). For factorized models, where the log probability can be written as a sum of terms involving only a single position, the mean dimension is *D_f_* = 1. For pairwise models, where the log probability contains terms depending on individual positions and position pairs, the mean dimension is 1 ≤ *D_f_* ≤ 2. For models that are dominated by higher-order interactions, we would expect a higher mean dimension. The mean dimension, therefore, allows us to assess the complexity of a model in terms of its interactions and, as we show in Methods, it can be estimated efficiently.

**Figure 1:**
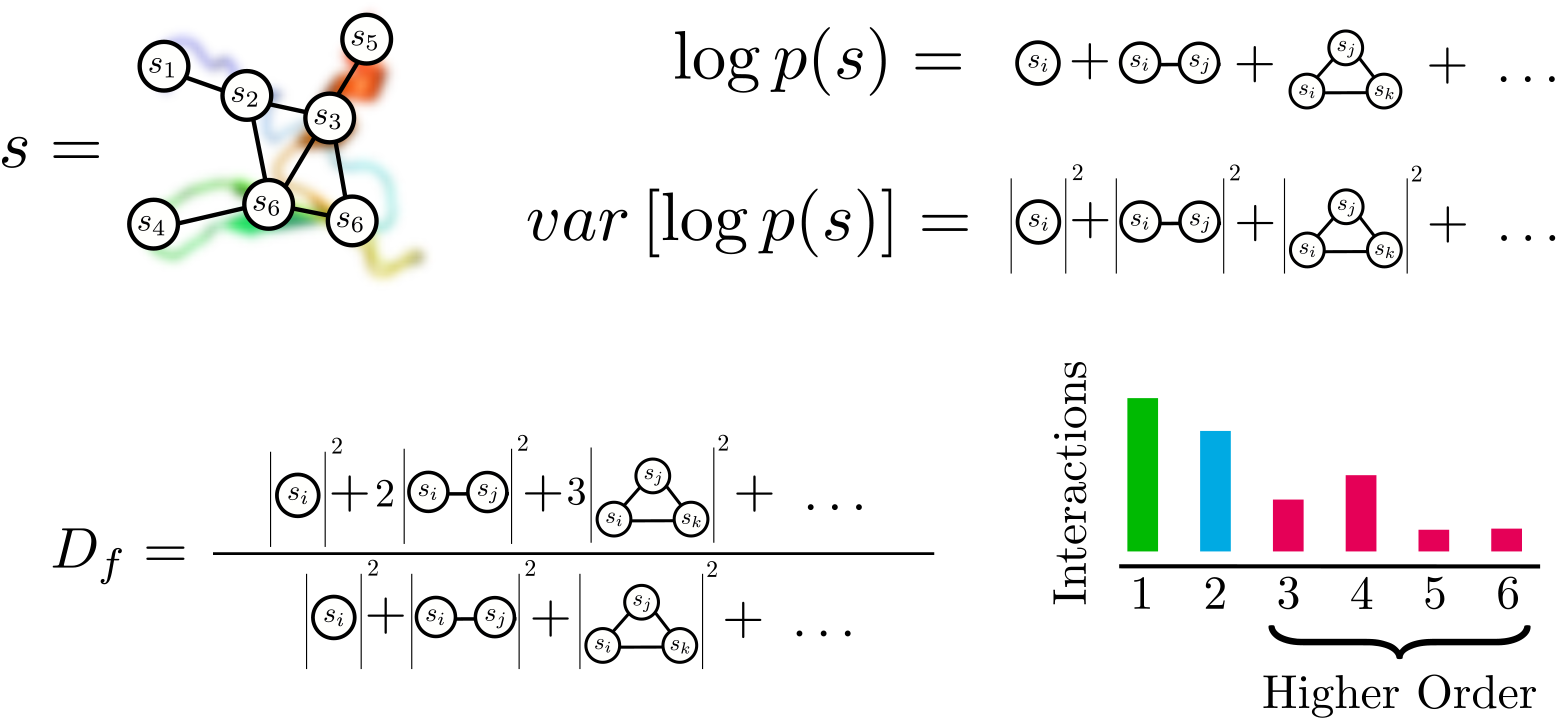
Overview: The figure represents a conceptual overview of the ideas used in this paper: The log probability log *p*(*s*) of a sequence *s* in a model can be expanded into terms of different orders. Under some assumptions on the expansion, the corresponding variance under the uniform distribution can be decomposed into contributions of different orders as well. The mean dimension is then defined as the average of orders under weights that correspond to contributions of orders to the total variance. The contribution of an order to the variance is proportional to the sum of squared interaction coefficients of that order. The mean dimension can therefore be conceptualized as an average over the histogram of squared interaction coefficients of different orders.

The mean dimension is a known tool in sensitivity analysis [19], and is also closely related to the *total influence* as defined for Boolean functions [22]. It has, however, to the best of our knowledge, never been applied to generative models for protein sequences. We derive the notion of mean dimension for such models in the framework of *energy-based models* (EBMs) of sequences of categorical variables in Methods. Given the specific framework and input domain, our derivation (see Methods) is different from typical approaches found in literature. To connect the present work to existing literature, we add in the Appendix a short review of the formalism as found in other works.

## 2 Methods

We assume a probabilistic, generative model that assigns a probability *p*(*s*) to sequences of amino acids *s* = (*s*_1_,…, *s_N_*) of fixed length *N*. Here, the sequence element si corresponds to a single amino acid or an alignment gap, and we will denote by *q* the number of possible values any *s_i_* can take (typically 20 different amino acids and a gap symbol, so *q* = 21). Such probability distributions underlie many popular generative models for protein sequences, for example Potts Models [9], Variational Autoencoders [1] or ArDCA [23].

### 2.1 Expansion

In this work, we will not analyse the probability distribution *p*(*s*) directly, but a function proportional to the negative log probability *f*(*s*) ~ – log *p*(*s*), where we assume that *p*(*s*) > 0 for all *s*. The function *f*(*s*) therefore maps sequences s of categorical variables of length *N* to a real value. We assume *q* to be the number of different categories, where a single category corresponds to an amino acid. Given *f*(*s*), the probability distribution *p*(*s*) can be recovered by

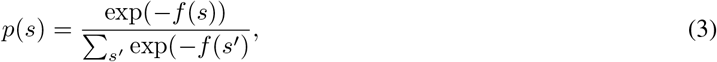

which is the Boltzmann-Gibbs distribution with *f*(*s*) as the energy. Here, the sum is over all possible sequences *s*′. We do not assume to be able to calculate the normalization factor corresponding to the denominator in Eq. 3, which is often not tractable, for example for Potts models.

A general expansion of *f* can be written as [17]

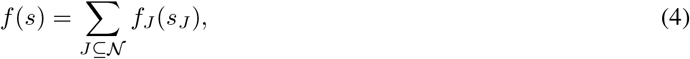

where we denote by 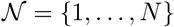 the set of all positions, by *J* a subset of 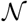, by *s_J_* the amino acids at positions in *J* and by *f_J_* a function that depends only on the amino acids at positions in *J*. The sum is therefore over the powerset of 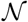.

While we can always expand a function *f*(*s*) in this way, see the Appendix, the expansion in Eq. 4 is in general not unique. For *L* ⊂ *J* we can for example redefine *f_J_*(*s_J_*) → *f_J_*(*s_J_*) + *f_L_*(*s_L_*) and *f_L_*(*s_L_*) → 0 without changing the values of *f*(*s*). Different ways of fixing these degrees of freedom are referred to as different *gauges* [11]. For the following, the choice of the *zero-sum* gauge is convenient, which is a popular choice for example for contact prediction [11]. In pairwise models, this gauge minimizes the Frobenius norm of pairwise interactions and leads to a representation in which as much as possible of the variance of *f*(*s*) is explained by simple terms depending only on a single position. In this sense, it aims to find the *simplest* representation of the expansion.

In the context of the expansion in Eq. 4 the zero-sum gauge, also called the *Ising gauge* [24], can be defined as the representation in which the sum of any *f_J_*(*s_J_*) over any of its dependencies is 0 for *J* ≠ ∅. This condition can be written as

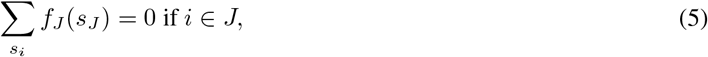

where we sum over all possible values of *s_i_* with the other dependencies of *f_J_* arbitrary. Every expansion can be written in this representation, which we show in the Appendix. We assume this representation from now on, if not stated otherwise. In this representation, it is easy to see that with 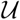 being the uniform distribution, where every amino acid in s is sampled independently with probability 1/*q*,

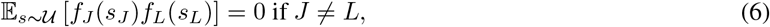

since the expectation, which contains a sum over all possible sequences, will contain a sum over a dependency of either *f_J_*(*a_J_*) or *f_L_*(*a_L_*) which is not a dependency of the other, resulting in 0 following Eq.5.

Similarly, the expectation of *f*(*s*) itself under 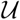 is

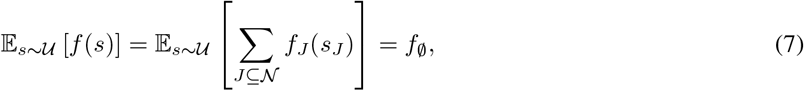

since the expectation for all terms except *f*_∅_ vanishes.

### 2.2 Mean Dimension for Generative Protein Sequence Models

We now define the mean dimension and begin by noting that the variance of *f*(*s*), when sampling *s* from the uniform distribution 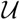, can be written as

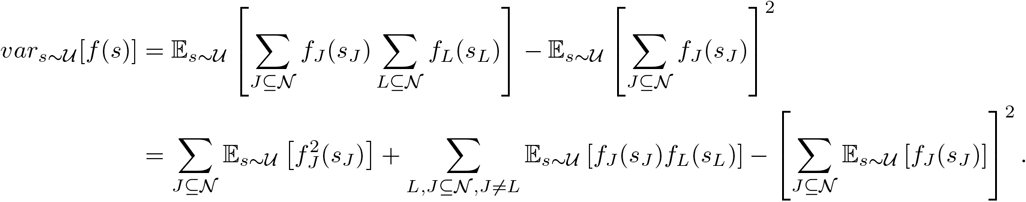

Assuming the zero-sum gauge defined by Eq. 5 and using the relations in Eq. 6 and Eq. 7, this leads to

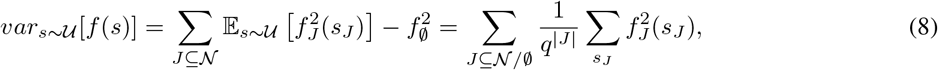

where |*J*| is the cardinality of *J* and the order of the interaction represented by *f_J_*. This can be written as a sum over interaction orders *k* as

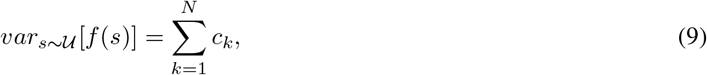

where we defined

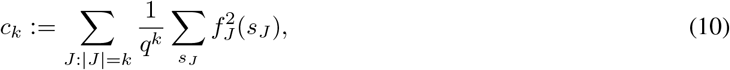

which is the contribution of order *k* to the variance, a sum over the averaged squared interactions coefficients corresponding to interactions of order *k*.

We can then define the weight *w_k_* of order *k* as

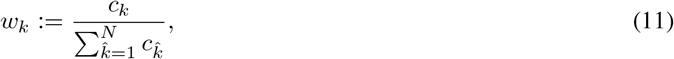

which satisfies *w_k_* ≥ 0 for all *k* and 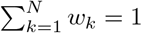. Note that the weights are only defined if 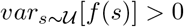, which we assume to be true in the following.

The mean dimension of interaction *D_f_* of the function *f*(*s*) can then be defined as the average of *k* with respect to these weights, leading to

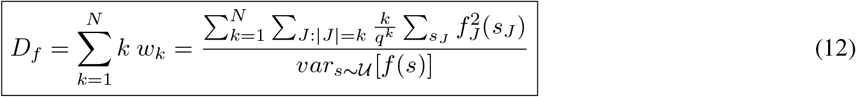

which is the average of interaction orders when weighted by their contribution to the variance of *f*(*s*) under the uniform distribution 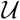. It is easy to see that *N* ≥ *D_f_* ≥ 1.

In the next section, we derive an efficient estimator for the mean dimension.

### 2.3 Estimating the Mean Dimension

We introduce the function 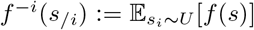, which is the function *f*(*s*) averaged over *s_i_* while keeping *s_/i_* fixed, where *s_/i_* is the sequence *s* without *s_i_*. When replacing *f*(*s*) with the expansion in Eq. 4 and using the definition of the zero-sum gauge in Eq. 5, we get

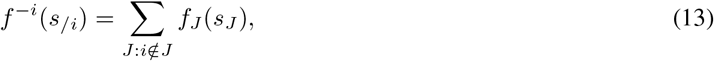

and, conversely,

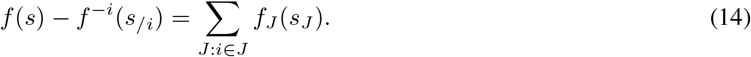

The same computation as for the variance above leads us to

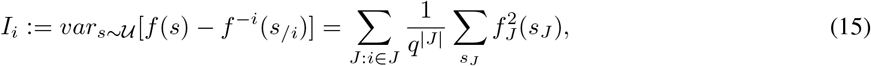

where *I_i_* can be interpreted as the part of the variance of *f*(*s*) that involves position *i*, also called the influence of *i* [22]. When summing the expression over *i*, a term involving the subset of positions *J* will appear exactly |*J*| times in the sum.

Therefore we can write

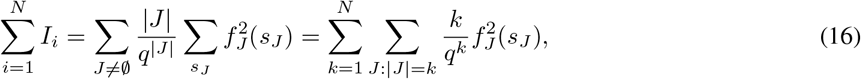

which is the numerator of the mean dimension in Eq. 12.

We finally arrive at

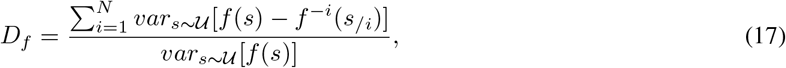

which can be approximated using samples from the uniform distribution 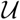. We will present a simple algorithm based on this identity in Methods.

## 3 Results

In the following we train different models on the MSAs of protein families and estimate their mean dimension, together with their performance on the task of mutational effect prediction. We take the data from [1], which contains 44 MSAs together with data from deep mutational scanning (DMS) experiments in the form of experimentally determined fitness changes following amino acid substitutions with respect to a wild-type protein. Since different experiments report different proxies for fitness, we follow the common approach and report Spearman correlation values of probabilities of mutated sequences in the models and the experimental measurements. We summarize the characteristics of the MSAs in the Appendix.

### 3.1 Estimating the Mean Dimension

The estimator in Eq. 17 can be directly translated into an algorithm for estimating the mean dimension of a given function *f*. We present a possible algorithm in Alg. 1, where we run for *T* iterations, sampling at every iteration a sequence from the uniform distribution for every position *i* and update based on these samples stream estimators tracking the quantities in Eq. 17. Since we need to calculate averages over all possible amino acids at all positions, which requires *q* evaluations of f at all *N* positions, a single iteration corresponds to *N* · *q* evaluations of *f* and the total number of necessary evaluations is *T* · *N* · *q*. Since every iteration can be seen as an independent estimate, we used bootstrap samples over the iterations to estimate the standard deviation of the final value. We found *T* = 1000 to be sufficient for reliable estimates for the protein families analyzed in the later sections.

We provide an implementation of this algorithm in the accompanying repository^2^. There we implement several further improvements and variations, for example parallelizing the loops or decoupling the sampling of sequences and the evaluation of *f* from the variance estimation, which allows for easier bootstrapping. The resulting estimator is, however, equivalent to Alg. 1.

#### Algorithm 1: Algorithm for Estimating the Mean Dimension *D_f_*

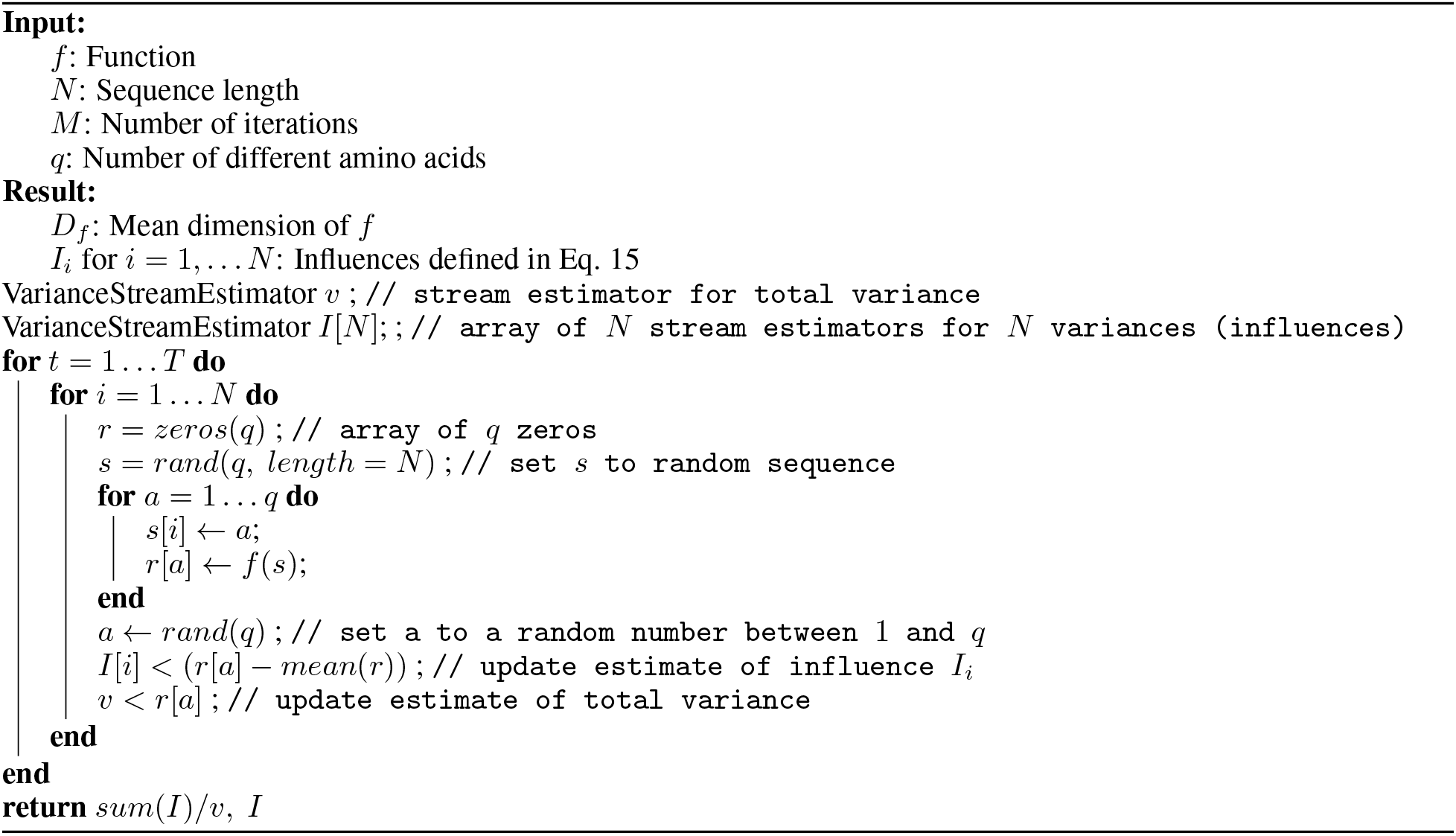

### 3.2 ArDCA

ArDCA [23] is a model for protein sequences that uses the autoregressive decomposition

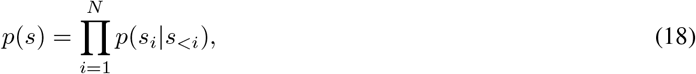

where *p*(*s_i_*|*s_<i_*) is the probability of the amino acid *s_i_* in position *i* given the part of the sequence that precedes position *i*, denoted by *s_<i_*. This probability is then defined as

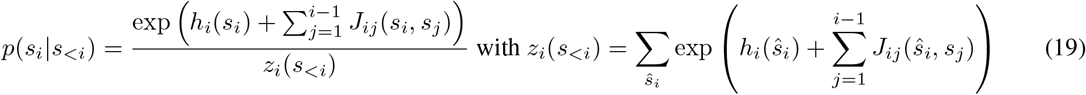

where *z_i_*(*s_<i_*) is a normalization factor depending on *s_<i_*. The terms *J_ij_*(*a, b*) are called couplings and depend on two positions, while the terms *h_i_*(*a*) are called fields and depend on a single position. Due to the autoregressive nature of the model, the probability of a sequence can be calculated directly using the equations above, which is an advantage to Potts models where the normalization constant is intractable and only a function proportional to the (log-)probability can be calculated. Note that due to the dependence of the normalization factors *z_i_*(*s_<i_*) on the preceding part of the sequence the model is not a pairwise model and can include higher-order interactions.

We use the code provided by the authors for training [23]. Training includes regularization using an *L*2 penalization with a different strength for the *J* and *h* terms, called *λ_J_* and *λ_h_* respectively. The authors provide different values optimized for generative modeling and mutational effect prediction. Since we will test the trained models on the latter task, we use the parameters optimized for mutational effect prediction (*λ_J_* = 0.01 and *λ_H_* = 0.0001) except where stated otherwise.

We retrained ArDCA on all 44 MSAs and estimated the mean dimension of the resulting models using *f*(*s*) = −log *p*(*s*). Surprisingly, we found all of the mean dimensions to be very close to 1, with an average of 1.02 and a maximal value of 1.053, see Table 1 in the Appendix. We suspected this to be due to the regularization constants used, where the couplings *J* have *a* regularization constant that is two orders of magnitude larger than the corresponding constant for the fields *h*.

From Eq. 19 it is clear that in the case *J* ≈ 0 the model factorizes and can be written as log 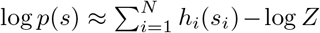, where *Z* does not depend on the sequence. Such a model would have indeed *D_f_* ≈ 1.

In order to test this hypothesis we retrained the models for a subset of four MSAs for BRCA1, GAL4, UBC9, SUMO1 with *λ_J_* varying logarithmically from 10^-6^ to 10^-1^. The resulting mean dimension estimates and the performance in terms of mutational effect prediction can be seen in Fig. 2. Several interesting results emerge: For a large range of λ_J_ the mean dimension for all models is relatively stable between 1.6 and 1.8, after which it rapidly drops to close to 1. Interestingly, this drop is very similar for all models and happens the roughly the same value of *λ_J_*. Furthermore, the *λ_J_* value at which the mean dimension for all models stabilizes towards 1, at *λ_J_* = 0.01, is exactly the value that the authors in [23] provided as optimal for mutational effect prediction. It seems therefore that, at least for mutational effect prediction, the best performing models are close to factorized models, where positions contribute almost independently to the log probability. Note, however, that this does not necessarily mean that a factorized model would perform equally to more complex models on the same data. The mean dimension is a measure taking into account *all* of the sequence space. Even if pairwise or higher-order interactions are comparatively small when viewed in this lens, they might still have a significant effect on the order of probabilities around a wild-type sequence.

**Figure 2:**
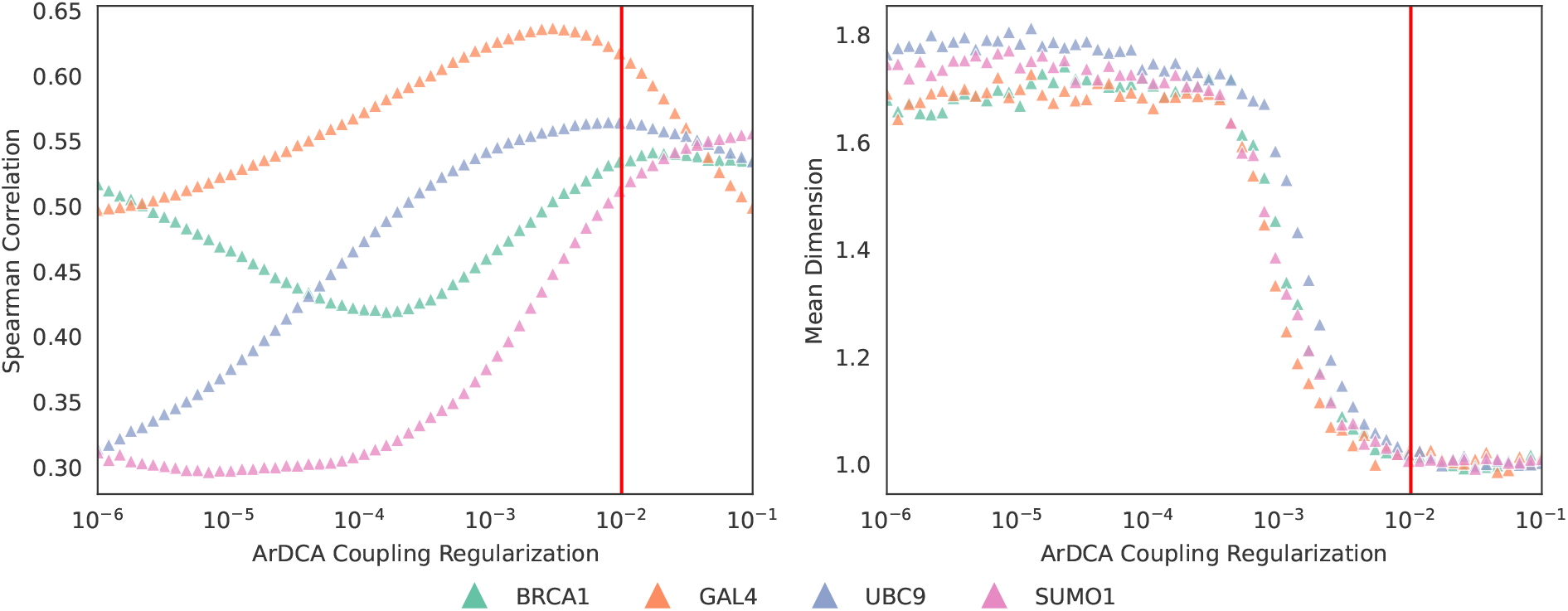
ArDCA Mean Dimension and Spearman Correlation. *Left*: The Spearman correlation with measured mutational effects for ArDCA models with varying *λ_J_* regularization. *Right*: Estimated mean dimension for the same ArDCA models. The red lines indicate the *λ_J_* regularization optimized for mutational effect prediction.

Next we analyzed the influence *I_i_* of positions as defined in Eq. 15, which measures the variance of *f* that involves position *i*. We show the influences in the right part of Fig. 3 for the ArDCA model trained on the GAL4 dataset. The influences vary significantly for different positions. We suspected that the influence of a position might be related to the evolutionary conservation of the position. We therefore calculated for every column in the training MSA the *site entropy* 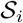, defined as

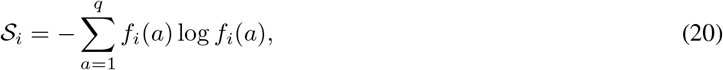

where *f_i_*(*a*) is the frequency of amino acid a in the consensus column *i* in the MSA. The higher the site entropy for a position, the more variable it is. In the left part of Fig. 3 we plot the site entropies of positions against their influences for the same ArDCA model as above. It is apparent that there is a strong *anti*-correlation between these two quantities. This can be explained by noting that a strongly conserved position in the training MSA will most likely lead to a model that assigns very low probabilities to sequences that contain an uncommon amino acid in this position. This, in turn, will lead this position to contribute significantly to the variance in log probability in sequences sampled from the uniform distribution, where all amino acids have equal probabilities in the position. By contrast, if a position is highly variable and has a large site entropy in the MSA, the model will most likely assign similar marginal probabilities to many different amino acids in this position. The position might still exhibit a large influence on the total variance, but this would need to be mediated by pairwise or higher-order interactions. These, however, are small for the ArDCA models used here, as shown above.

**Figure 3:**
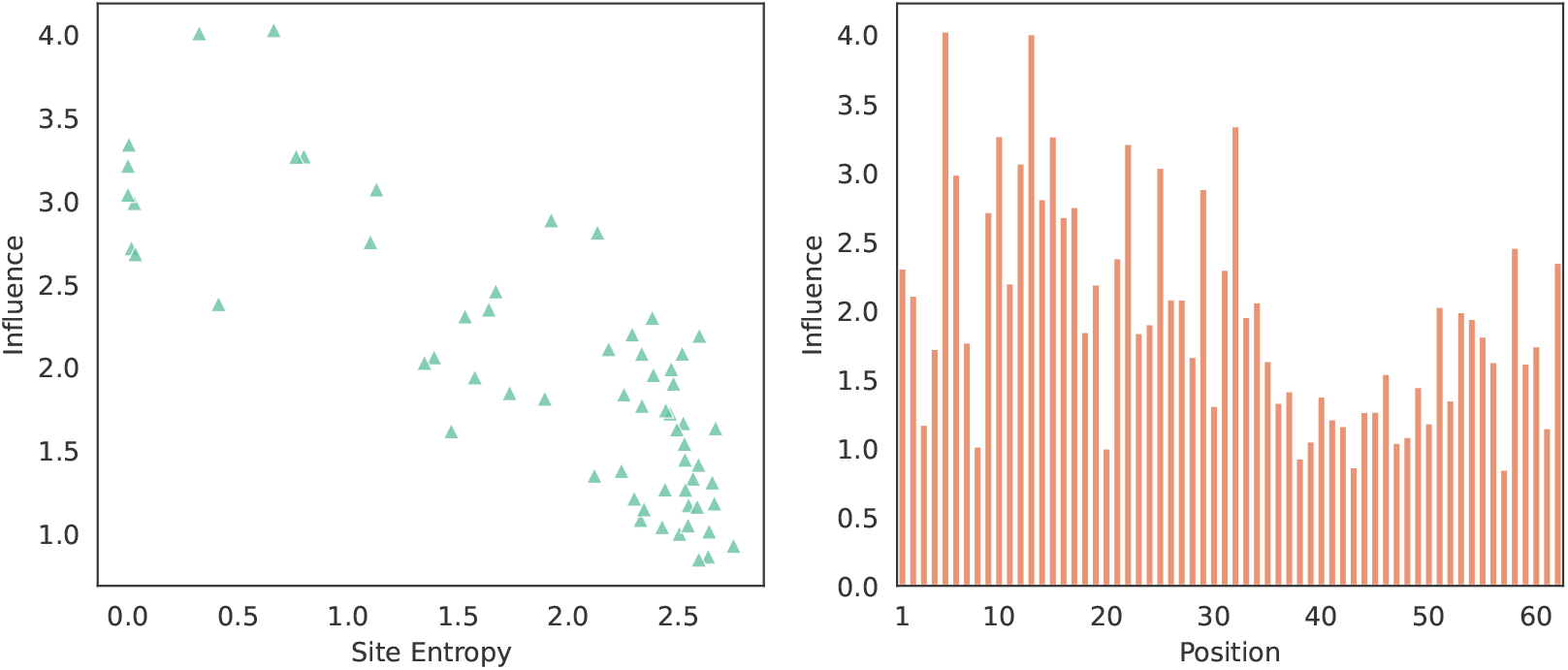
Site Entropies and Estimated Influences for ArDCA trained on GAL4. *Left*: Site entropy of a position in the training dataset versus the estimated influence of the position in the model as calculated from Eq. 15. *Right:* Influence for individual positions along the sequence. Both parts use the same model, trained with *λ_J_* = 0.01 and *λ_h_* = 0.0001.

### 3.3 Variational Autoencoder

Variational autoencoders (VAEs) are popular latent-space models for homologous protein data that have been shown to perform well in tasks like mutational effect prediction [1], the assessment of the pathogenicity of amino acid variants in humans [4], and the exploration of sequence space by analyzing the latent representations [13].

Following the VAE architecture presented in [13], we project the sequence s into a probability distribution over latent representation 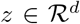 by using a probabilistic encoder MLP which takes a one-hot encoded version of *s* as input and outputs the means and log variances of a multivariate Gaussian with zero off-diagonal correlations. A second MLP, the probabilistic decoder, then transforms the latent representation to the inputs of a *sofimax* layer, which defines probabilities of amino acids at all positions. Defining the prior over *z*, the model can be trained using the ELBO objective [25]. The probability of a sequence *s* can then be estimated using importance sampling [26]. We use a single-layer MLP with 40 hidden units and a *tank* activation function for both the encoder and decoder, and a latent representation with dimension *d* = 5, with an *L*2 penalization of strength 0.01 on all parameters. These values perform well in practice on mutational effect prediction. Following the setting in [13], we train in full batch mode for 10.000 epochs.

For calculating the log probabilities necessary for determining the mean dimension we use 5000 ELBO samples, which also leads to a good performance in mutational effect prediction. However, this means that a single evaluation of *f* in the estimator in Eq. 17 requires 5000 single evaluations of the neural network. We therefore do not estimate the mean dimension on all 44 MSAs but on the same subset that we used for a more focused analysis in the last section (the MSAs corresponding to BRCA1, GAL4, UBC9 and SUMO1). We calculate the mean dimension of the final models and, in order to get more insight about the training process, during training. We also monitor the Spearman correlation with the experimental data during the same checkpoints. We show both quantities for the four protein families in Fig. 4.

**Figure 4:**
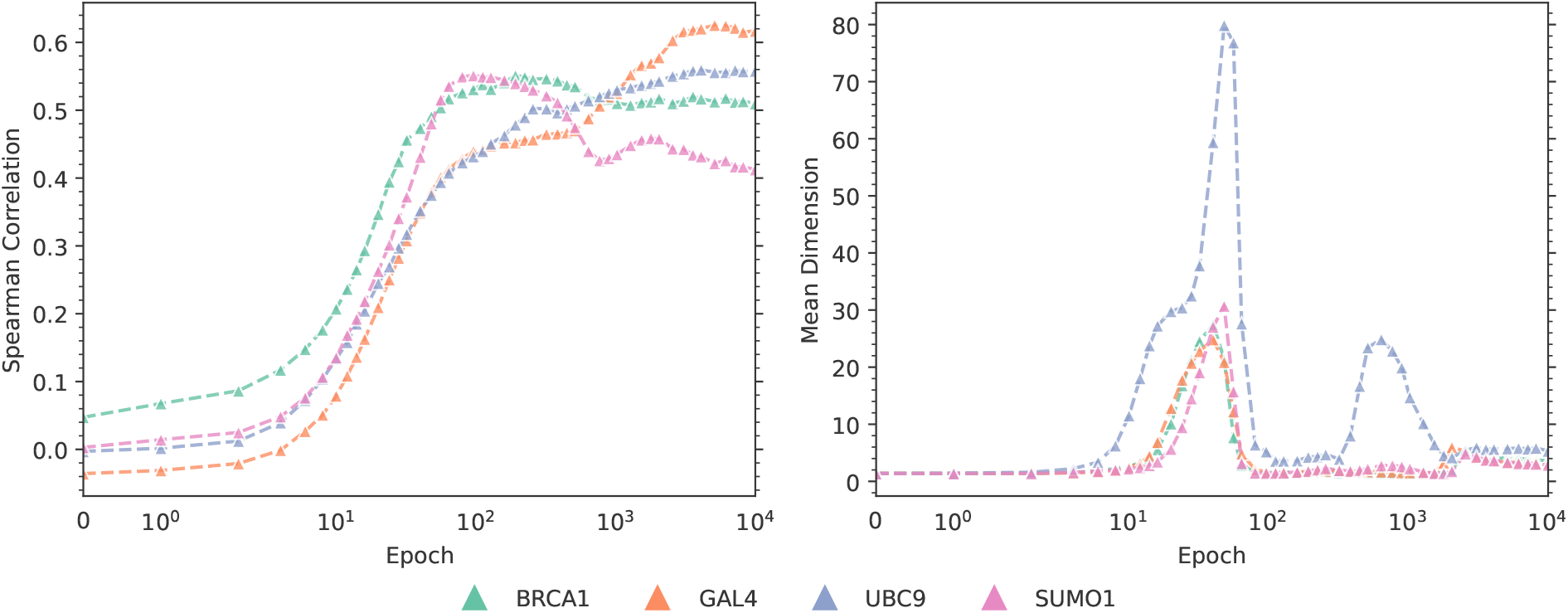
Spearman Correlation with Experimental Mutational Data and Mean Dimension during Training for VAE models. *Left:* Spearman correlation with the experimental fitness effects for different protein families during training. *Right:* Estimated mean dimension for different protein families during training.

Focusing on the first 250 epochs, the behavior of the mean dimension is similar for all datasets: The initial mean dimension is close to 1 at epoch 0 (corresponding to the models with randomly initialized parameters), but then increases in the first 50 epochs to around 20 for BRCA1, GAL4 and SUMO1 and to about 80 for UBC9. This is interesting since it means that the models acquire strong higher-order components during the first phase of training. However, in the next 50 epochs the mean dimension decreases again to close to the initial values. In this first phase, the Spearman correlation with the experimental mutational data increases rapidly until about epoch 100. In a second phase, from about epoch 100 to 3000, the mean dimension remains relatively low for BRCA1, GAL4 and SUMO1, but shows another peak of height around 30 for UBC9. The Spearman correlation with the experimental mutational data fluctuates in this phase, for example increasing for BRCA1 to a peak and then decreasing again. After about epoch 4000 both the mean dimension and the Spearman correlation stabilizes. For SUM01, the Spearman correlation seems to decrease slightly until the end of training. We note that this MSA is particular as factorized models (so models with no pairwise or higher-order interactions and with mean dimension equal to 1) perform *better* than more complex models in terms of mutational effect prediction [1]. The final mean dimensions are significantly higher than the corresponding values for ArDCA, ranging from 2.8 for SUMO1 to 5.3 for UBC9, see Table 1 in the Appendix. This seems to indicate that the VAE models indeed implement higher-order interactions that are important for explaining the variance on sequences from the uniform distribution, in contrast to the ArDCA models. Since their performance in terms of mutational effect prediction is not better than for ArDCA models (compare Fig. 2), these higher-order interactions do not seem to be important for this task, however.

We also show the site entropy and the influences in Fig. 5. The situation is quite different than for ArDCA models in that there is no discernible correlation between the influence of a position and its site entropy in the MSA. In addition, the influences vary less among the positions. This is surprising since it indicates that all positions contribute equally to the model variance, independent of their conservation in the training set. In order to analyse this effect close we also calculate the variance in log probability when exchanging amino acids at a given position while keeping the rest of sequence fixed to a wildtype sequence from the training set. More precisely, we calculate the quantities

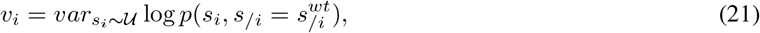

for all positions *i*, where 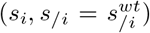 is a shorthand for the wildtype sequence *s^wt^* with 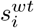 replaced by *s_i_* and the variance is taken over *s_i_* following the uniform distribution 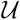. Using the yeast sequence for GAL4 we show the quantities and their relation to the site entropy in Fig. 6. The variance in log probability is correlated with the site entropy and the distribution of this quantitiy along the sequence is very similar to the one obtained from ArDCA, see Fig. 3. Together, these results indicate that the influences of the positions are similar to each other when looking at the global distribution, but differentiate for parts of the sequence space closer the training set. We note that this is in line with the idea that VAEs trained on protein sequences behave like mixtures of factorized models, where the emitted factorized model depends on the coarse location in sequence space [16]. Such models might exhibit a large mean dimension due to the mixture, but have a very simple structure locally.

**Figure 5:**
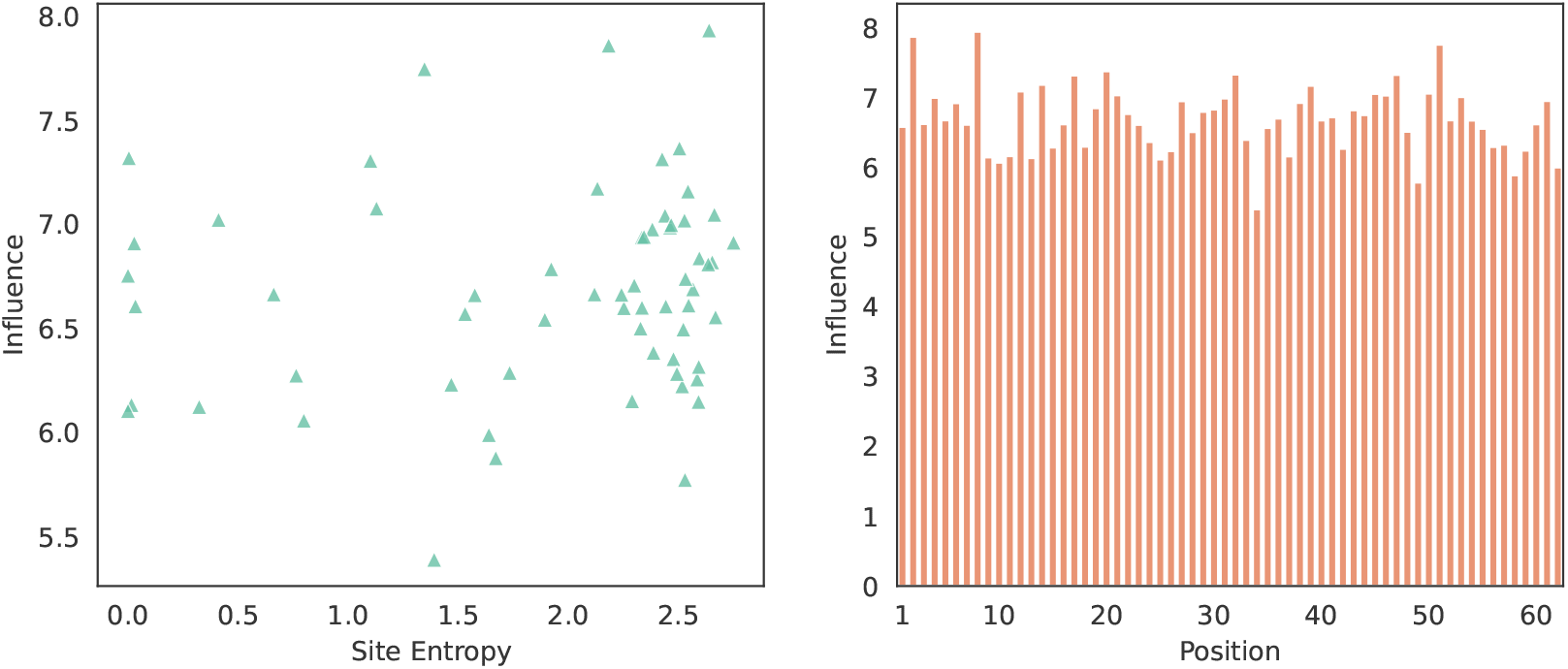
Site Entropies and Estimated Influences for VAE trained on GAL4. *Left:* Site entropy of a position in the training dataset versus the estimated influence of the position in the model as calculated from Eq.15. *Right:* Influence for individual positions along the sequence.

**Figure 6:**
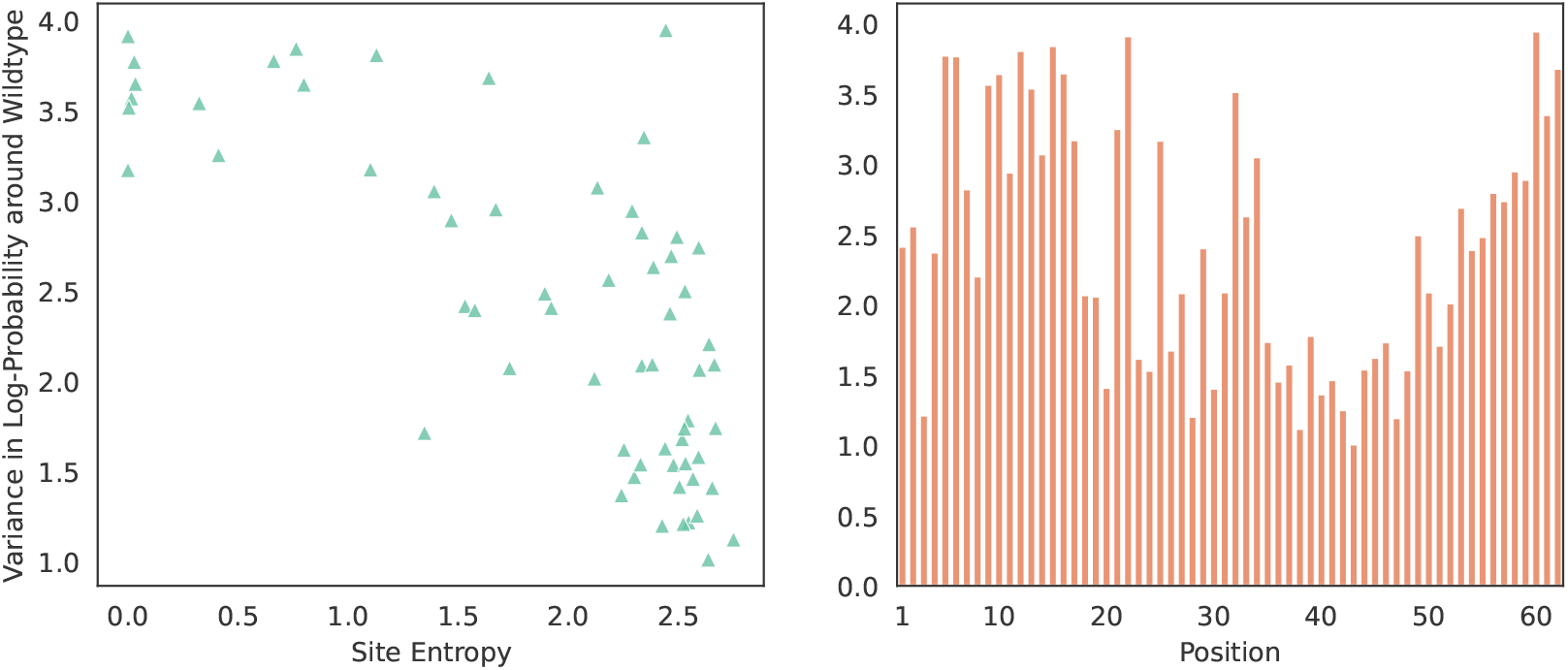
Site Entropies and Variance in Log Probability around Wildtype for VAE trained on GAL4. *Left:* Site entropy of a position in the training dataset versus the variance in log probability as calculated from Eq. 21. *Right:* Variance in log probability around wild type for individual positions along the sequence.

## 4 Discussion

In this work, we have introduced the mean dimension of interaction for generative protein sequence models, which measures the average order of interaction coefficients in a general expansion of the log probability assigned by the model to protein sequences. We derived a simple estimator of the mean dimension and an algorithm for the computation. We estimated the mean dimension of two model classes, ArDCA and the VAE, trained on MSAs of different protein families. We analyzed changes in mean dimension when modifying the regularization constants in ArDCA and during training for the VAE. The results indicate that the mean dimension of ArDCA models is close to 1 when using parameters optimized for mutational effect prediction and we never observed a mean dimension larger than 2 for this model class in any setting. For the VAE, the mean dimension is higher and we also showed it to be highly variable during training. We also highlighted that the mean dimension is related to the performance of the models in terms of mutational effect prediction.

We believe the mean dimension to be an interesting analysis tool in that it can be efficiently calculated while having a clear interpretation in the context of higher-order interactions, which are often cited as a central advantage of complex models trained on protein sequence data.

The main limitation of the mean dimension presented here is that it is defined via a decomposition of the variance over the whole sequence space. The average dimension of important interactions when assigning the same weight to all possible sequences might, however, be very different from the average dimension of important interactions for example on the training set. A model with a low mean dimension could for example still include subtle higher-order interactions that are important in a specific region in sequence space. This is closely connected to the question of different gauge representations, which makes the concept of interactions itself ambiguous, since their strength depends on the representation used. Fixing a gauge representation, as we did in this work, removes this freedom, but other choices for the representation are possible and will lead, in general, to different results. A possible avenue could be for example to ask for the representation that leads to the least complex expression on the training set, and then measure the mean dimension in this representation. We intend to explore these questions in further research.

While we used the mean dimension in this work for model comparison and analysis, it would be interesting to probe its potential as an element of the model selection or training procedure itself, for example as an indicator for early stopping or during hyperparameter optimization. In addition, it is possible that the mean dimension could reveal further insights into other phenomena observed in neural networks, for example the double descent phenomenon.

## A Appendix

### A.1 Dataset and Mean Dimension

**Table A1:**
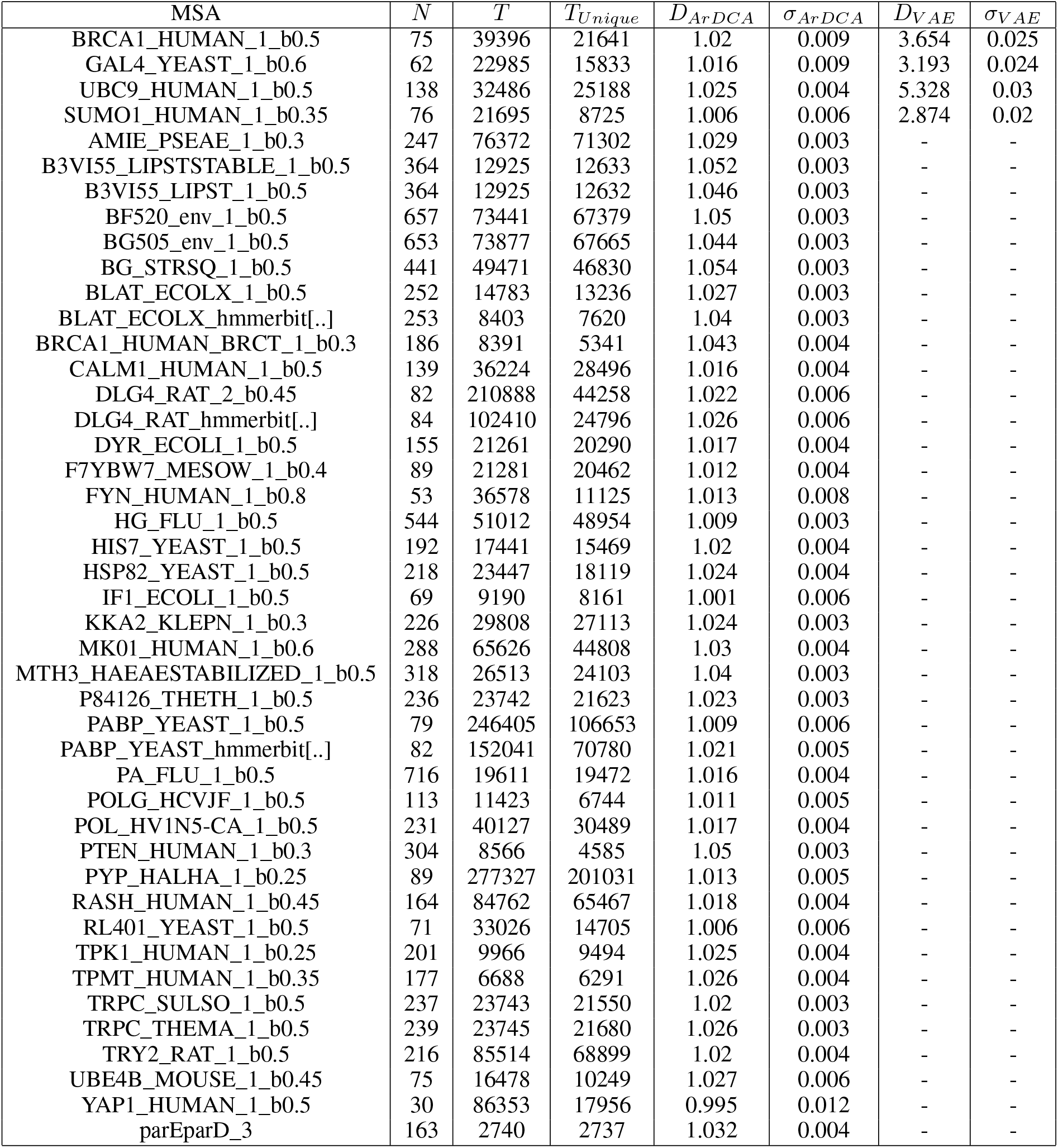
MSA Properties and Mean Dimension. The table gives an overview over the MSAs used, their properties and the estimated mean dimension. The columns are: ***MSA***: MSA filename in the supplementary data in [1]; ***N***: Sequence length (only consensus columns); *T*: Number of sequences in the MSA; *M_Unique_*: Number of unique sequences in the MSA; ***D*_{*ArDCA,V AE*}_**: Mean dimension estimated for ArDCA and VAE models; ***σ*_{*ArDCA,V AE*}_** Standard deviation of mean dimension estimates using 1000 bootstrap samples.

### A.2 Review of Mean Dimension

The mean (or effective) dimension of a function 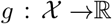, denoted with *D_g_*, is typically defined as the average size of interactions present in a multivariate input-output mapping (we reserve the notation *f* for the special case of functions defined on categorical inputs and use *D_f_* for their mean dimension). Intuitively, consider an function mapping a *d*-dimensional input 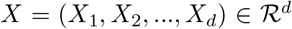 to a real value 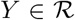, where we consider the inputs following some cumulative distribution *F_X_*. In machine learning applications, this could be for example the log probability of a class in a neural network trained for image classification. Works such as [27, 28, 29] show that it is possible to expand the input-output mapping in the analogue of a multi-body expansion as the sum of 2^d^ component functions, of which *d* are univariate component functions, each depending only on a single *X_i_*, 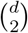 are bivariate component functions, each depending only on a single pair (*X_i_*, *X_j_*), and so on. If the expansion contains only univariate components, then the mapping is additive (it is the sum of univariate functions in each variable) and the mean dimension is equal to 1. Conversely, a value of the mean dimension greater than 1 indicates that interactions are present. Given that the expansion parameters are defined using only the inputs and outputs, without further knowledge about how one is calculated from the other, the mean dimension is an ideal tool for exploring the complexity (or richness) of black-box input-output mappings. Recently, renewed interest in the use of the mean dimension in machine learning was generated [20, 21]. The key idea of these works is to use the mean dimension as a tool for comparing different neural network architectures. For instance, [21] consider fitting several well known convolutional neural network architectures of different sizes to the same image dataset. Their experiments show that the mean dimension assumes values much smaller than the overall problem dimensionality *d*, and that it correlates with properties like the depth of the neural networks. The same finding is confirmed in the experiments of [20].

From the technical side, the works of [20] and [21] are able to avoid a technical complication: Following directly the definition of mean dimension, its computation would be infeasible in most practical applications since it contains an exponential number of terms. However, [20] and [21] show that, assuming feature independence, it is possible to compute the mean dimension using the expression

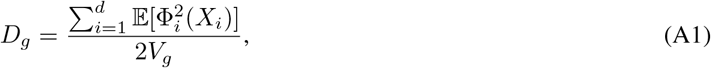

where *V_g_* is an estimate of the variance of the target *Y* if the inputs follow the distribution *F_X_*, and, for each feature, and the random variables 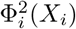 are defined as 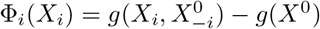, where *X*^0^ follows the distribution *F_X_* and 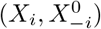 represents *X*^0^ with 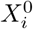 exchanged with a newly sampled *X_i_*. We derive a very similar expression for our specific case in the methods section of the main paper.

Note that the mean dimension introduced in this section depends on the function under consideration and the distribution of inputs. The case where features are not independent under this distribution presents a major challenge for the calculation of the mean dimension, and the identity in Eq. A1 does not hold anymore. The result of replacing the distributions with, e.g., uniform distributions is hard to interpret in general and heuristic approaches have to be introduced, for example the introduction of de-correlating PCA layers into analysed neural networks [21].

Fortunately, for protein sequences the situation is different: As we show in the methods section of the main paper, basing our estimates on the uniform distribution over amino acids in the sequences leads to an estimate for the mean dimension that can be interpreted as the mean dimension defined via the parameters in a specific *gauge representation* of the log probability, where different gauge representations refer to different way of fixing degrees of freedom in the expansion of the log probability. The gauge representation corresponding to our values of the mean dimension is the so-called *zero-sum* or *Ising* gauge, which is widely in use in the field [6, 11, 17].

### A.3 Zero-Sum Gauge

In the following we show that we can represent every real-valued function defined over fixed-sized amino acid sequences in the zero-sum gauge. This proof is already outlined in [17], and also mentioned in [30], but we present a more detailed version here since it is central to the present work. Note that some quantities are defined differently from there, which we will point out below.

Formally, we assume 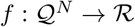 to be a function which maps a sequence of *N* categorical variables 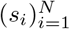 to a real number, where the variables take values from the finite set 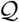 with *q*:= |*Q*|.

We then show that we can write the function as

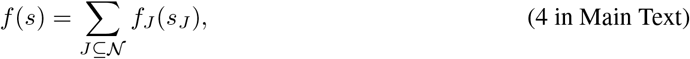

where 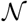 is the set of all positions and the *f_J_*(*s_J_*), which are functions depending only on the subset of positions 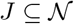, satisfy the conditions

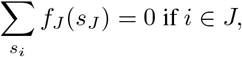

defining the so-called *zero-sum* gauge, which is also sometimes called the *Ising* gauge.

We first notice that we can expand any function *f* trivially by defining

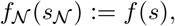

with *f_J_*(*s_J_*):= 0 for |*J*| < *N*. This representation corresponds to assigning the function values of *f* on all *q^N^* sequences to the *q^N^* interactions of order *N* and setting all other interactions to 0. This shows that some representation of the form in Eq. 4 can always be found.

In the following we do not assume *f_J_*(*s_J_*) = 0 for |*J*| < *N* but that we are given some representation with interactions at arbitrary orders.

Given this representation, we define transformed interactions 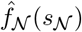 of order *N* as

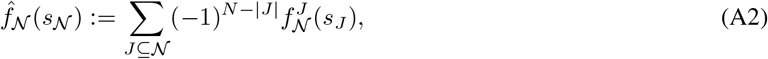

where we define

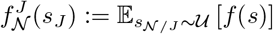

as the function *f*(*s*) after averaging out *s_N/J_*, with *s_N/J_* following the uniform distribution 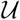. This leaves a function that depends on the positions in *J*. Note that 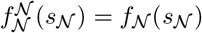. We also point out that the notation here is different from [17], where 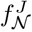 is defined as a function depending on all positions *except* the ones in *J*. For the current purpose, the notation used here is clearer, however.

We now define transformed interactions for lower orders than *N* as

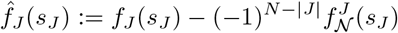

for all 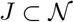.

The value of the function 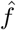 for a given sequence *s*, defined by the transformed interactions, is then

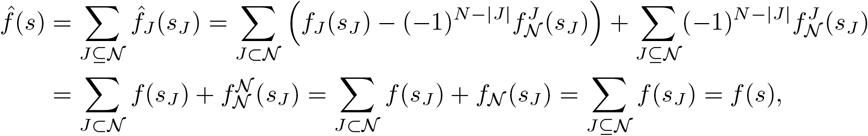

so the transformation leaves the function value invariant.

Furthermore, if we sum the transformed interactions of order *N* over any dependency *s_i_*, we get

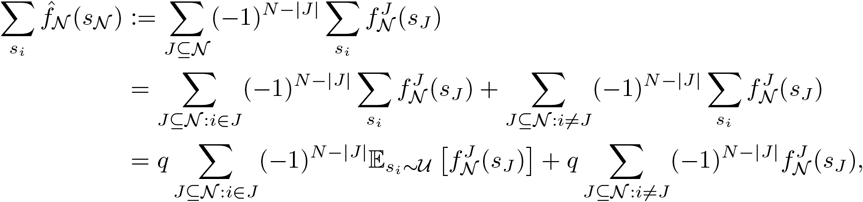

where we separated the sum over *J* into terms that depend on *s_i_* and terms that do not depend on *s_i_* in the first line and replaced the sum over *s_i_* with an expectation over *s_i_* following the uniform distribution 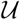 in the second line. From the definition in Eq. A2 it can be seen that the term 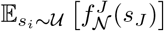 is identical to 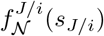, since we just add another position to the expectation.

The sum in the first term can therefore be rewritten as

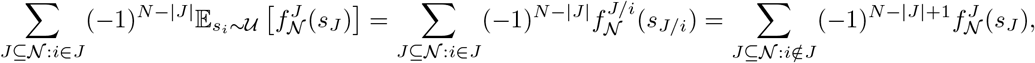

which leads finally to

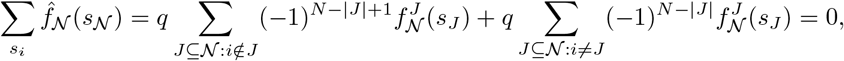

which means that the interactions at order *N* in the transformed representation satisfy the zero-sum conditions outlined in the main paper.

Fixing the interactions at order *N*, the problem is reduced to transforming a model with interactions up to order *N* – 1 to the zero-sum gauge. This, however, can be achieved by the same procedure starting at order *N* – 1. We note that it is not trivial to give an expression for the final parameters at all orders, but following the procedure, starting at order *N* and continuing until order 1, will certainly transform the original representation into the zero-sum gauge.

## Footnotes

2 https://github.com/christophfeinauer/ProteinMeanDimension

